# Development of a long-read NGS workflow for improved characterization of fastidious respiratory mycoplasmas

**DOI:** 10.1101/2022.03.31.486658

**Authors:** Isaac Framst, Cassandra D’Andrea, Monica Baquero, Grazieli Maboni

## Abstract

Mycoplasmas are respiratory pathogens in humans and animals and due to their fastidious nature, they have been historically underdiagnosed. Lack of standardised diagnostic, typing and antimicrobial susceptibility testing methods makes clinical management and epidemiological studies challenging. The aim of this study was to develop a cost-effective and accurate sequencing workflow for genotypic characterization of clinical isolates of respiratory mycoplasmas using a rapid long-read sequencing platform. Critical aspects of bacterial whole genome sequencing were explored using fastidious respiratory *Mycoplasma* (*M. felis* and *M. cynos*) isolated from animals including: (i) four solid and liquid-based media based on a specialized formulation for *Mycoplasma* culture, (ii) three DNA extraction methods modified for sequencing purposes, and (iii) two *de novo* assembly platforms as key components of a bioinformatic pipeline including Flye and Canu assemblers. DNA quality and quantity compatible with long-read sequencing requirements were obtained with culture volumes of 160ml in modified Hayflick’s broth incubated for 96 hours. The other three culture approaches investigated did not meet the DNA quality criteria required for long-read sequencing. The use of bead-beating bacterial cell lysis in the extraction protocol resulted in smaller fragments and shorter reads compared to enzymatic lysis methods. Overall, Flye generated more contiguous assemblies than the Canu assembler. This novel study provides a step-by-step sequencing workflow including mycoplasma culture, DNA extraction and *de novo* assembly approaches for the characterization of highly fastidious respiratory mycoplasmas. This workflow will provide diagnosticians, epidemiologists, and researchers with a more comprehensive tool than the laborious conventional methods for a complete genomic characterization of respiratory mycoplasmas.

## 1 Introduction

Mycoplasmas are often implicated in human and animal respiratory diseases as primary or secondary pathogens ^1–3^. Due to their reduced genome and dearth of many biosynthetic pathways, these fastidious bacteria depend on rich medium for growth leading to the lack of standardised diagnostic, typing and antimicrobial susceptibility testing (AST) methods. This makes clinical management and epidemiological studies of mycoplasma-associated respiratory infections challenging. The presence of several *Mycoplasma* species in clinical samples may also complicate diagnostic procedures ^4–6^. As there are more than 100 different species among pathogenic and commensal mycoplasmas, species identification is critical for a comprehensive characterization of respiratory infections ^7,8^. Current AST and *Mycoplasma* spp. identification at species level require culture-based and molecular methods, which takes two to three weeks ^8^. This is considered unsuitable for routine diagnostic purposes, even in specialized laboratories ^6,8^. An alternative to AST is the identification of genotypic changes within loci associated with phenotypic resistance and treatment failure ^6^. Common molecular approaches for *Mycoplasma* identification include PCR and Sanger sequencing of the 16S rRNA gene and intergenic spacer region (ISR) ^5,7^ These assays often have low specificity in the presence of multiple *Mycoplasma* species ^4^. Further strain typing of mycoplasmas using multi locus sequence typing requires additional PCR assays, sequencing or DNA microarrays targeting specific loci, increasing diagnostic complexity ^9,10^.

Currently available next-generation sequencing (NGS) technologies may aid to overcome some of diagnostic issues of mycoplasmas. With the various advantages and decreasing costs of NGS, the accurate genotypic characterization could complement or even replace the culture-based phenotypic tests. Long-read DNA sequencing technologies, such as that offered by Oxford Nanopore Technologies (ONT), are promising new tools for fast, cost-effective, and high throughput whole genome sequencing (WGS), which can be applied to fastidious respiratory bacteria ^11,12^. Long-reads may allow for more efficient *de novo* assembly over short-read by decreasing computational requirements, leading to faster and more complete assemblies ^11,13^.

Despite the considerable potential of NGS, several challenges must be addressed. There are currently no standardized approaches that provide reliable procedures for NGS of fastidious mycoplasmas. Thus, this study aimed to develop a step-by-step workflow for long-read WGS of mycoplasmas to provide diagnosticians, epidemiologists, and researchers a better tool for rapid identification, *in silico* AST, virulence and strain typing. The development of a robust WGS workflow will fill the need for more accessible and complete genomic characterization of *Mycoplasma-associated* infections.

## 2. Methods

### 2.1 Study Design

Different approaches for long-read Nanopore sequencing of *Mycoplasma cynos* and *Mycoplasma felis* were assessed in this study, including four culture methods, three DNA extraction methods, and two genome assembly pipelines. The three *M. cynos* and two *M. felis* strains included in this study were isolated from the bronchoalveolar lavage of dogs and cats, respectively, as part of the diagnostic workup completed by the Animal Health Laboratory (AHL), University of Guelph, between 2016 and 2018 (Table 1). The archived isolates were provided by AHL and kept at −80°C in a modified Hayflick’s broth with 10% glycerol until analysis.

**Table 1.** Mycoplasma cynos and *Mycoplasma felis* strains used in this study.

| Species | Isolate number<sup>a</sup> /name | Source | Cell viability<sup>b</sup> |
| --- | --- | --- | --- |
| *M. cynos* | 18-030510-1/cynos 1 | Canine Bronchoalveolar Lavage | 69.9 % |
| *M. cynos* | 16-078726-2/ cynos 2 | Canine Bronchoalveolar Lavage | 70.6 % |
| *M. cynos* | 17-098999-3/ cynos 3 | Canine Bronchoalveolar Lavage | 68.9 % |
| *M. felis* | 16-040612-4/ felis 4 | Feline Bronchoalveolar Lavage | 79.0 % |
| *M. felis* | 16-057152-4/ felis 6 | Feline Bronchoalveolar Lavage | 69.8 % |

<sup>a</sup> Isolates provided by the Animal Health Laboratory, University of Guelph.
<sup>b</sup> Percent viability measured for 160 ml cultures after 96-hour incubation period (culture approach 4) using flow cytometry. See Methods section for further details of culture approach 4.

### 2.2 Preparation of *Mycoplasma* cultures for DNA extraction

Four different approaches to obtain *Mycoplasma* cells for DNA extraction were assessed: 1) scraping of colonies from the agar surface; 2) agar plugs containing *Mycoplasma* colonies on top; 3) 40 ml of modified Hayflick’s broth incubated for 48 h; and 4) 160 ml of modified Hayflick’s broth incubated for 96 h (see supplementary file 1 for agar formulation). A considerably smaller culture broth volume was initially attempted (6 ml); however, the concentration of DNA obtained was too low and for this reason this approach was excluded from this study (see supplementary file 4 for unsuccessful culture and DNA extraction methods).

For <u>approach 1</u>, *M. felis* and *M. cynos* strains were retrieved from −80 °C stocks and plated on 50 mm plates containing 10 ml modified Hayflick’s agar, and incubated at 37 °C, 5% CO_2_ for 48 h. Then, the entire surface of the agar plate was scraped using a loop and deposited into 180 μl of lysis solution provided by the GenElute Bacterial Genomic DNA extraction kit (Sigma-Aldrich, Darmstadt, Germany). For <u>approach 2</u>, *Mycoplasma* cultures on agar were prepared as described in approach 1, then three agar plugs were excised from the plate using a disposable loop to pierce the agar creating a ‘plug’ of about 1 cm^2^. The area of the plate containing the highest observed colony density was chosen for plug excisions. Once excised, plugs were placed in 180 μl of lysis solution, with no pelleting steps. For <u>approach 3</u>, flasks containing 40 ml modified Hayflick’s broth were incubated at 37 °C, 5% CO2 for 48 h in a vented culture flask (see supplementary file 2 for broth formulation). For assessment of approach 4, the initial culture volume (40 mL) was increased to 160 ml after 48 h of incubation and kept for an additional 48 h in the incubator, totalling to 96 h prior to extraction. DNA extractions were conducted immediately following incubation for all four approaches.

Pelleting of *Mycoplasma* cells from broth cultures was performed using 50 ml tubes with a high-speed Fiberlite™ centrifuge rotor (Thermo Scientific). For small broth cultures, one tube was used; for large cultures, broth was distributed equally between three tubes, then combined into a single tube after pelleting. Samples were centrifuged for 45 minutes at 14,000 x g at ambient temperature. Approximately 1.5 ml of supernatant was retained in the tube to resuspend the pellet using a wide bore pipette. Gentle scraping of the tube side walls was necessary for complete resuspension. The resuspended pellet was transferred into a 2 ml microcentrifuge tube for re-pelleting at 16,000 x g for 10 minutes. The supernatant was decanted until approximately 100 μl remained covering the pellet, which was then resuspended as the initial material for DNA extraction.

### 2.3 Viability of *Mycoplasma* cells in modified Hayflick’s broth

Flow cytometry was conducted on all five isolates using BD FACS Accuri C6 (Becton Dickinson). Isolates were prepared using culture approach 4. Samples for flow cytometry were prepared and analyzed using a previously described method ^14^. Flow cytometry data was analyzed for cell viability using FlowJo v. 10.7.2 ^14^

### 2.4 Assessing three different methods for *Mycoplasma* DNA extraction

For DNA extraction, modifications were made to the manufacturer’s protocol, including elution buffer substitutions, elution temperature and column incubation time as described below.

<u>Extraction method 1</u> used a modified protocol with the GenElute Bacterial genomic DNA kit (Sigma-Aldrich). The pellet was resuspended in 180 μl of kit-included lysis buffer. To prevent residual RNA from interfering with downstream processes, 20 μl kit-included 10 mg/ml RNase A was added prior to the initial lysis incubation period. Samples underwent enzymatic lysis for 30 minutes at 55 °C without orbital shaking. Final elution was standardized across all DNA extraction methods with a volume of 100 μl in non-EDTA 10 mM tris-HCL elution buffer EDTA is known to interfere with downstream sequencing (Omega Bio-Tek, Norcross USA) (pH 8.5 and at 70 °C) ^15^. The elution buffer was applied to the spin column and incubated for five minutes at room temperature before centrifugation. Extraction method 1 was performed for samples from culture approaches 1, 2, 3 and 4.

<u>Extraction method 2</u> used a modified protocol for the Omega Bio-tek E.Z.N.A. tissue kit (Omega Bio-tek, Norcross USA) due to its practicality and low cost (CAD $1.14/sample). Following initial pelleting as described above, pellets were lysed mechanically with low binding zirconium beads (100 mg) (OPS Diagnostics, Lebanon USA) using a Qiagen TissueLyser II for four minutes at 30 Hz. Samples were then briefly centrifuged to collect beads at the bottom of the tube, followed by enzymatic lysis for 1 h at 55 °C as described in the manufacturers protocol. To eliminate residual 20 μl 10 mg/ml RNase A was added prior to the initial lysis incubation period.Final elution was performed as described in method 1. DNA extraction method 2 was performed for samples from culture approaches 3 (small broth) and 4 (large broth).

<u>Extraction method 3</u> used a modified version of the Qiagen QIAamp Mini DNA kit (Qiagen, Venlo, Netherlands). Following initial pelleting, and mechanical lysis (as described in extraction method 2), samples were briefly centrifuged to collect beads at the bottom of the tube. The manufacturers protocol was then followed, with 1 h of enzymatic lysis time. Final elution was performed using the previously described method. Method 3 was performed for samples from culture approaches 3 (small) and 4 (large broth).

### 2.5 DNA quantification and quality assessment

DNA quantification and quality were assessed using nanodrop 1000 photospectrometer (Thermofisher) and Qubit 2.0 (Thermofisher). Qubit working solution was prepared using Qubit dsDNA BR assay according to the manufacture’s recommendations (Thermofisher, Waltham USA). Fragmentation analysis was performed using Agilent TapeStation 4150 (Agilent, Santa Clara, USA) according to the manufacturer’s recommendations.

### 2.6 ONT library preparation and sequencing

A total of 11 DNA samples representing the different DNA extraction methods were chosen for WGS. Library preparation was performed using the Rapid Barcode kit RBK004 (Oxford Nanopore Technologies, Oxford, UK) following the manufacturer’s directions. Total DNA input was adjusted to 400 ng prior to pooling of samples. The sequencing run was performed with minION mk1B with minKNOW v.21.06.0 software.

### 2.7 Whole genome assembly and quality assessment

All raw reads were submitted to a genomics pipeline developed in this study (Figure 1). Basecalling was performed post-run using High Accuracy (HAC) model GPU basecalling with automatic GPU compute resource allocation in GUPPY basecaller v5.0.12 (Oxford Nanopore Technologies). Reads were demultiplexed and trimmed using Porechop v0.2.4 (https://github.com/rrwick/Porechop). Genomes were assembled using Flye v2.8.3, with an additional parallel pipeline using Canu v2.2 to investigate the effect of genome assembly method on genome quality for *Mycoplasma* (https://github.com/marbl/canu). Potential base-call errors in the draft assembly were verified and corrected by Racon v.1.4.20. Minimap2 v2.21 (https://github.com/lh3/minimap2)^16^ was used to generate overlaps, consensus, and read-assembly maps for Racon. Assembly quality statistics were generated using QUAST v5.0.2 (http://quast.sourceforge.net/index.html) ^17^. Rapid genome annotation was conducted using Prokka v1.14.6 (https://github.com/tseemann/prokka) ^18^.

**Figure 1.**
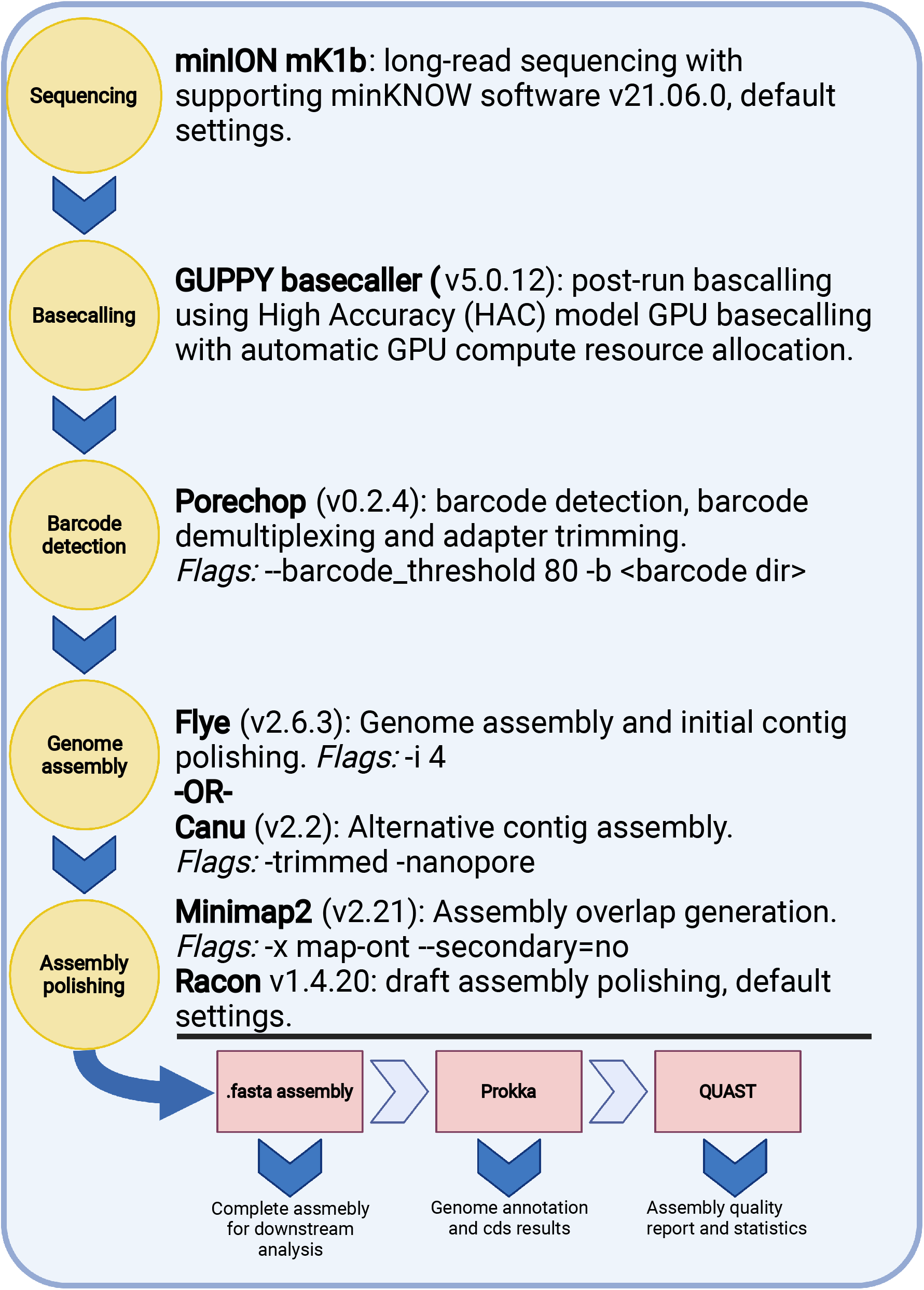
Bioinformatic pipeline for *de novo* assembly of long-read sequencing of *Mycoplasma cynos* and *Mycoplasma felis* generated by Oxford Nanopore Technologies.

The statistical analysis including t-tests and one way analysis of variance (Kruskal-wallis) were performed using R statistical software (v. 1.2.1335) ^19^.

## 3 Results

### 3.1 Acquisition of *Mycoplasma* cells for DNA extraction

All archived isolates remained viable after thawing from −80°C in a 10% glycerol solution, and they remained viable after 96 h of incubation as demonstrated by flow-cytometry (Table 1).

Initial attempts to culture *Mycoplasma* for DNA extraction were unsuccessful. *Mycoplasma* cells obtained using the agar scraping and agar plug approaches did not yield significant DNA concentrations. The 6 ml broth culture produced detectable but low concentrations which did not meet the requirements of long-read sequencing. A list of DNA extraction results with low quantity and purity along with their corresponding culture conditions is provided in Supplementary file 4.

All three DNA extraction methods showed acceptable DNA purity for large and small broth cultures. However, extraction method 1 yielded purer DNA than the other two methods (i.e. average is closer to 1.8 for 260/280, and 2.2 for 260/230 ratios) (Table 2).

**Table 2.**
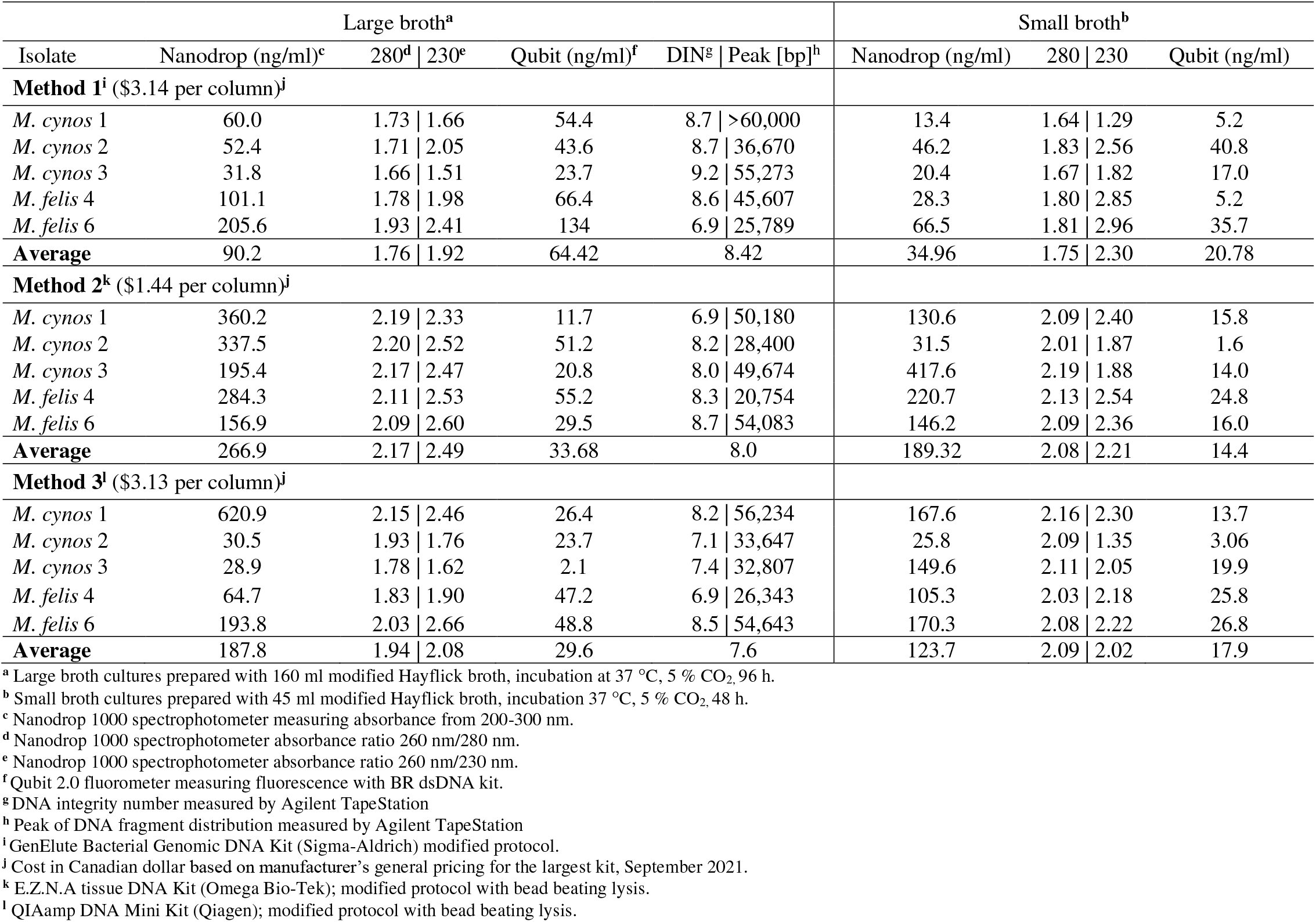
DNA extraction performance of three different extraction methods using two different *Mycoplasma* culture approaches.

Regarding DNA concentration, large broth cultures (culture method 4) yielded significantly higher DNA concentrations compared to small broths (culture method 3) (Table 2). Thus, large broth extractions were chosen for downstream library preparation and DNA sequencing (Table 2). When comparing the three DNA extraction methods, all produced acceptable DNA concentrations with large broth cultures, except for *M. cynos* 3, method 3 (Table 2).

DNA fragmentation analysis was performed for all large broth extractions. Most of the isolates presented high DNA integrity number (DIN > 8) (11/15), with extraction method 3 having the lowest average DNA integrity (Table 2). This study also investigated the use of bead-beating lysis on downstream ONT analysis. For DNA extraction methods 2 and 3, bead-beating lysis in combination with enzymatic lysis were used, while method 1 only used enzymatic lysis. Based on the peak fragment size (bp), methods using the bead-beating lysis produced shorter fragments compared to the enzymatic-only lysis method (extraction method 1) (Table 2).

### 3.2 Assembly statistics

We recovered genome lengths consistent with the expected genome sizes in each of the three DNA extraction approaches as shown by QUAST analysis. Most isolates were successfully assembled into a single contig, except for *M. felis*-4 being assembled into 4 and 10 contigs using extraction methods 2 and 3, respectively (Table 3). The high N50’s, low L50’s and low N’s indicated contiguous assemblies across kits and assembly methods except for *M. felis*-4 extraction methods 2 and 3 (Table 3). There was no significant difference regarding the average read length between extraction methods, as extraction method 1 resulted in an average read length of 4,434 bp (*s* = 2,043 bp), while method 2 of 3,573 bp (*s* = 1189 bp), and method 3 of 3,114 bp (*s* = 11,06 bp) (Table 3). All kits generated sequencing coverage greater than 40x using Flye assembler ^20^, but extraction method 1 produced a higher average coverage compared to 2 and 3 (Table 3).

**Table 3.** Sequence run and assembly statistics using Flye and Canu assemblers.

| Isolate | Average read length [bp] | no. Reads | Coverage<sup>a</sup> | No. contigs \| Total length [bp] |  | N50 \| L50<sup>b</sup> |  |
| --- | --- | --- | --- | --- | --- | --- | --- |
|  |  |  |  | Flye | Canu | Flye | Canu |
| Method 1<sup>c</sup> |  |  |  |  |  |  |  |
| *M. cynos* 1 | 5,263 | 65,593 | 363x | 1 \| 951,262 | 3 \| 1,058,580 | 951,262 \| 1 | 983,043 \| 1 |
| *M. cynos* 2 | 5,144 | 42,219 | 228x | 1 \| 951,622 | 1 \| 985,908 | 951,622 \| 1 | 985,908 \| 1 |
| *M. cynos* 3 | 6, 971 | 47,661 | 329x | 1 \| 1,009,272 | 1 \| 1,034,656 | 1,009,272 \| 1 | 1,034,656 \| 1 |
| *M. felis* 4 | 3,013 | 54,843 | 167x | 1 \| 988,020 | 3 \| 1,043,847 | 988,020 \| 1 | 620,867 \| 1 |
| *M. felis* 6 | 1,778 | 150,644 | 268x | 1 \| 945,799 | 1 \| 958,423 | 945,799 \| 1 | 958,423 \| 1 |
| Method 2<sup>d</sup> |  |  |  |  |  |  |  |
| *M. cynos* 2 | 3,045 | 20,410 | 65x | 1 \| 950,900 | 2 \| 980,529 | 950,900 \| 1 | 961,155 \| 1 |
| *M. cynos* 3 | 5,220 | 23,464 | 121x | 1 \| 1,008,899 | 1 \| 1,025,417 | 1,008,899 \| 1 | 1,025,417 \| 1 |
| *M. felis* 4 | 2,455 | 56,944 | 144x | 4 \| 973,334 | 2 \| 985,836 | 551,770 \| 1 | 728,442 \| 1 |
| Method 3<sup>e</sup> |  |  |  |  |  |  |  |
| *M. cynos* 2 | 2,903 | 13,233 | 40x | 1 \| 950,735 | 2 \| 966,046 | 950,735 \| 1 | 949,977 \| 1 |
| *M. cynos* 3 | 4,311 | 66,246 | 283x | 1 \| 1,008,981 | 1 \| 1,033,760 | 1,008,981 \| 1 | 1,033,760 \| 1 |
| *M. felis* 4 | 2,129 | 19,096 | 41x | 10 \| 945,789 | 3 \| 973,826 | 255,270 \| 2 | 347,844 \| 2 |

<sup>a</sup> Coverage calculation method: total reads [bp] / assembled genome size [bp]. <sup>b</sup> N50 is the length of the shortest contig at half of the total genome length; L50 is the smallest number of contigs composing half of the total genome size. <sup>c</sup> GenElute Bacterial Genomic DNA Kit (Sigma-Aldrich); modified protocol. <sup>d</sup> E.Z.N.A tissue DNA kit (Omega Bio-Tek); modified protocol with bead beating lysis. <sup>e</sup> QIAamp DNA mini kit (Qiagen); modified protocol with bead beating lysis.

Both, Flye and Canu assemblers produced high quality assemblies, but Flye successfully assembled 9 of 11 isolates into a single contig, whereas Canu was successful in only 5 of 11 isolates (Table 3). PROKKA annotation counts indicate similar performance of both assemblers (supplementary file 5), as there was no significant difference in annotation counts between assembly methods (*t*)13 = 1.43, *p* = 0.17.

A summary of the resulting recommended step-by-step workflow from culture to assembly analysis is illustrated in Figure 2.

**Figure 2.**
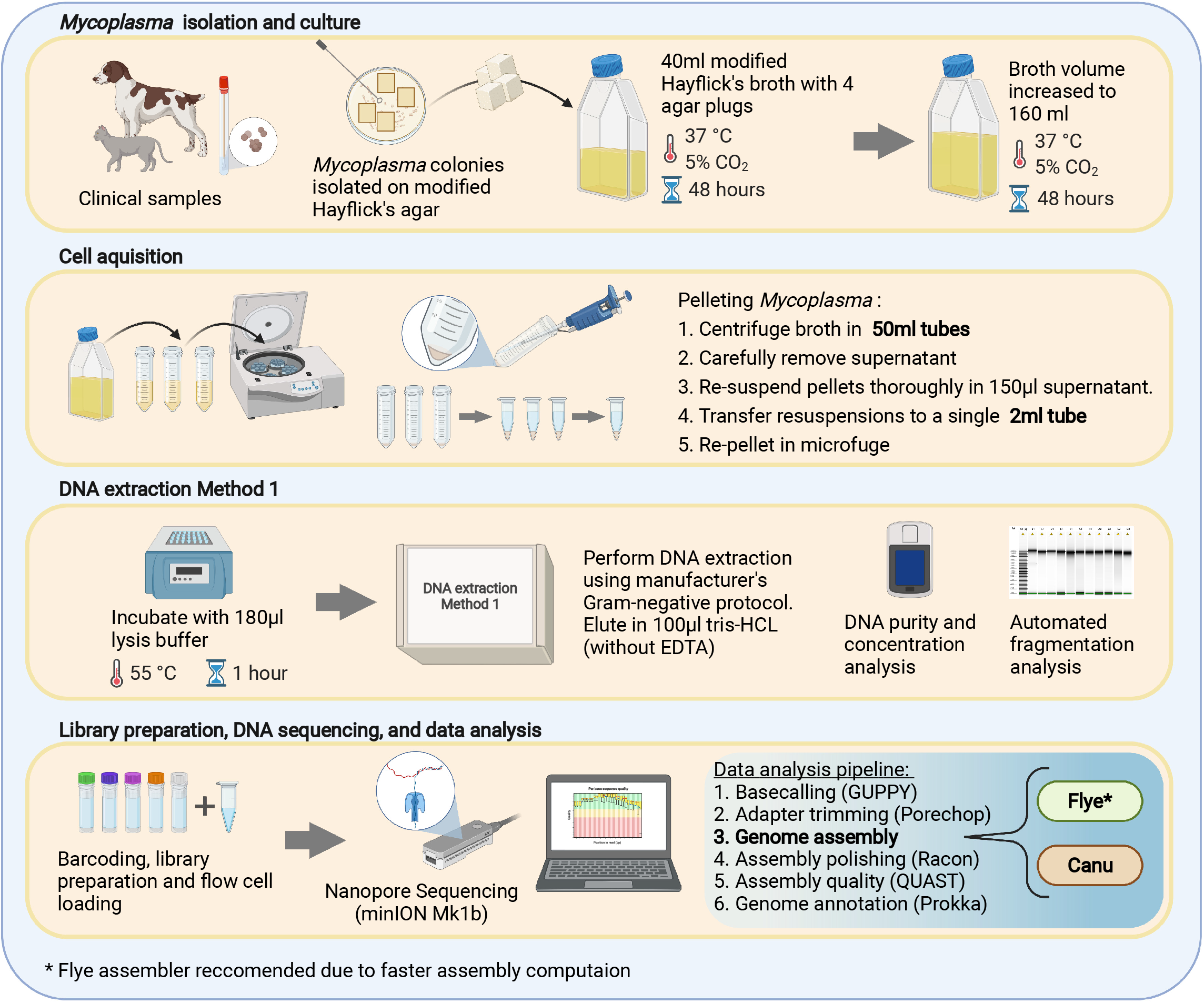
Workflow for whole genome sequencing of respiratory mycoplasmas using Oxford Nanopore Technologies long-read platform. The illustration highlights only the approaches and methods which presented the greatest performance in this study.

## 4 Discussion

We assessed a step-by-step workflow, with multiple alternatives for each step, for long-read NGS of respiratory mycoplasmas to develop an optimal standard procedure. Three critical aspects were explored: (i) solid and liquid-based medias using a specialized formulation for *Mycoplasma* culture, (ii) three DNA extraction methods modified for sequencing compatibility, and (iii) two *de novo* assembly platforms as key components of a bioinformatic pipeline.

We initially investigated several culture approaches for acquisition of *Mycoplasma* cells aiming high DNA concentrations as required for long-read sequencing. Agar plate scraping and agar plugs yielded low or undetectable DNA concentrations. These findings were inconsistent with a previous study that obtained enough DNA of *M. cynos* from agar plugs using the same DNA extraction kit ^21^. Therefore, our next attempt was to grow mycoplasmas in broth cultures (6 mL) incubated overnight, but DNA quantities were below ONT recommendations, likely because of slow growth and small volume of broth (supplementary file 4). This contrasts with other species of mycoplasmas, such as *M. bovis* and *M. arginini*, which grow well in 8 mL of liquid culture ^14^. As different *Mycoplasma* species have distinct growth rates, this might be associated with faster growth of *M. bovis* and *M arginini* than *M. cynos* and *M. felis* ^14,22^. We have observed that *M. felis* strains grow more abundantly than *M. cynos* on solid media (data not shown). As in 40 mL cultures (small broths) DNA concentrations were below ONT recommendation, we had to increase the broth volume to 160 mL and lengthened incubation from 48 to 96 hours. The alternative of culturing the isolates in one step with a large volume of broth (160 mL) for 96h of incubation did not support enough *Mycoplasma* growth. Therefore, growing the isolates in two steps from a small volume (40 mL) followed by a larger volume (160 mL) supported more bacterial growth and delivered the highest quantity and quality of DNA.

All three DNA extraction methods produced enough data with ONT sequencing (Table 3). We observed that the bead-beating lysis methods 2 and 3 resulted in more DNA fragmentation and lower read lengths. As mycoplasmas lack cell wall, mechanical lysis methods such as bead-beating might not be needed. Regarding assemblies, only extraction method 1 produced single-contigs. Therefore, we observed that the choice of DNA extraction method may influence sequencing performance (Table 3). The literature evaluating the potential impact of DNA extraction workflows on subsequent whole genome sequencing of mycoplasmas is limited if not absent. For other bacteria such as *E. coli*, solid-phase extraction kits, as the ones investigated in this study, were preferred over salting-out kits for sequencing purposes ^15,21^. Based on performance and user experience, we recommend the solid-phase DNA extraction method 1 for long-read NGS-based workflow of respiratory mycoplasmas.

A previous study examining a variety of DNA quantification methods for NGS found that Nanodrop photospectrometer measured higher DNA concentrations than expected, being unreliable at low DNA concentrations. In comparison, Qubit estimations were more accurate ^23^. We also observed considerable discrepancies between both measurements which is in accordance with previous studies ^23^. Therefore, estimations based only on Nanodrop could incorrectly identify that the DNA concentration is sufficient for ONT sequencing.

Many *de-novo* assemblers exist for long-read sequencing, however, not all assemblers are suitable for genomes with low GC content or highly repetitive regions like mycoplasmas. High coverage and read length are important to generate contiguous assemblies, especially in the presence of extensive repetitive regions; however, high coverage is not guaranteed when working with fastidious mycoplasmas where it may be difficult to obtain high quality DNA extracts for sequencing ^13,20^. As this bacterium has a very small genome, less overall sequence data might be needed for good coverage, possibly explaining why we obtained complete genome assemblies even for samples with input DNA concentrations below ONT recommendations. Interestingly, there were two instances where both assemblers failed to generate single-contig assemblies despite ≥40x coverage recommended for Flye. Likely, these instances indicate uneven sequencing depth at specific loci or sample contamination. The Flye assembler successfully assembled more isolates into a single contig than Canu but predicted gene numbers were similar for both assemblers. In addition, Flye used less computational memory compared to Canu for the same input data, which improved the overall assembly time and computational requirements of the pipeline. Benchmarking data of long-read assemblers for bacterial whole genome assemblers suggest that better circularization, contiguity, and shorter runtime is generally achieved using Flye, especially for lower coverage applications ^24,25^. Our results and experience are consistent with these benchmarks.

A limitation of this study is that financial constraints allowed for only one ONT sequencing run, which resulted in a lower number of replicates than desired.

## 5 Conclusion

A WGS workflow for long-read sequencing of fastidious respiratory mycoplasmas was developed in this study. *M. cynos* and *M. felis* were used in the optimizations, but considering their slow growth and fastidious nature, we speculate that this workflow could also be applied to other highly fastidious respiratory mycoplasma species. Our findings indicate that large liquid cultures, 160 mL Hayflick’s broth, yielded best results for sequencing, and DNA extraction method 1 provided the best user experience and quality metrics performance. We observed that the more cost-effective method 2 provided sufficient DNA input for ONT sequencing but with a low coverage, and depending on the purpose of the downstream analysis, this method could still be satisfactory. Overall, Flye generated more contiguous assemblies than the Canu assembler. To our knowledge, this is a novel study to provide a comparison of different approaches for a long-read WGS workflow of respiratory mycoplasmas. This type of genetic based tool is increasingly needed for species identification, strain typing and selection of proper antibiotic therapy in *Mycoplasma*-associated respiratory infections.

## Acknowledgements

We would like to thank Jutta Hammermueller and Dr. Ksenia Vulikh for laboratory technical support, and Dr. Patrick Boerlin from the University of Guelph for constructive criticism of this manuscript.

## Data Availability

Bioinformatic pipeline shell script with dependencies, install instructions and usage instructions is available at https://github.com/iframst/FRM_assm.

Sequences generated as part of this study are available from the National Center for Biotechnology Information (NCBI) genome database (https://www.ncbi.nlm.nih.gov/genome) under accession number SUB11225409. Individual genome accession numbers are listed in Supplementary file 3.

## Supplementary files legend

**Supplementary file 1**. Instructions to prepare Hayflick’s modified agar. Proportions for 1000 ml.

**Supplementary file 2**. Instructions to prepare modified Hayflick’s broth.

**Supplementary file 3**. GenBank Accession numbers for the whole genome sequences of *Mycoplasma cynos* and *Mycoplasma felis* generated in this study. GenBank accession numbers for assembled genomes.

**Supplementary file 4.** Different approaches used for bacterial culture, cell acquisition, and DNA extraction that were unsuccessful in this study.

**Supplementary file 5**. Genome annotations of *Mycoplasma cynos* and *Mycoplasma felis* using PROKKA.

